# Sex Differences in Cortical Morphometry during Ageing: Examining the Interplay between Lifestyle and Reproductive Factors

**DOI:** 10.1101/2021.10.14.464259

**Authors:** Manuela Costantino, Aurélie Bussy, Grace Pigeau, Nadia Blostein, Gabriel A. Devenyi, Ross D. Markello, Raihaan Patel, Nicole Gervais, M. Mallar Chakravarty

**Affiliations:** Cerebral Imaging Centre, Douglas Mental Health University Institute, Verdun, Canada; Undergraduate program in Neuroscience, McGill University, Montreal, Canada; Integrated Program in Neuroscience, McGill University, Montreal, Quebec, Canada; Department of Biological and Biomedical Engineering, McGill University, Montreal, Canada; Department of Psychiatry, McGill University, Montreal, Canada; Rotman Research Institute at Baycrest, Toronto, Ontario, Canada; Department of Psychology, University of Toronto, Toronto, Ontario, Canada

**Keywords:** cortical thickness, magnetic resonance imaging, menopause, partial least squares, pregnancy

## Abstract

Sex differences in neurodegenerative disorder prevalence have been attributed to life expectancy, modifiable risk factors related to lifestyle and the impact of changes in sex hormones and the reproductive system. Although these factors are known to interact with one another, they are often studied in isolation. Here, we used a multivariate approach to investigate how lifestyle, along with menopause and the number of children, interacts with cortical thickness (CT) in healthy adults. Using CT measures from T1-weighted scans (MPRAGE, 1 mm^3^ voxels; 124 participants; 67 females; 40-70 years old) from the Cam-CAN dataset. Using a partial least squares decomposition, we identified patterns of covariance between CT and lifestyle factors, menopause and the number of children. In women, we identified significant patterns that linked education, socioeconomic status, social contact and length of reproductive period to CT in the left prefrontal cortex, as well as alcohol consumption, physical activity and menopausal status to CT in the frontal poles. Contrastingly, the results in men were driven by education and anxiety, and involved increased CT in the temporal poles. Our findings suggests that sex differences in cortical anatomy during brain ageing might be driven by interactions between contrasting lifestyles and the female-specific endocrine environment.

## Introduction

As the global population ages and the incidence of neurodegenerative disease rises, increased attention has been brought to the study of the brain during older age (Peters 2006). However, given the continued search for effective treatments, there is a clear need to characterize potential risk factors for maladaptive ageing. In order to identify biomarkers of disease, a detailed characterization of the interaction between biological and environmental factors involved in healthy ageing can help to understand their effects on brain anatomy and behaviour (Barnes and Yaffe 2011; Norton et al. 2014;Livingston et al. 2017).

Neuroimaging techniques have largely contributed to this goal by allowing researchers to study the anatomy of the brain in very large numbers, using non-invasive procedures. Cortical thickness (CT) measures serve as a useful proxy for the anatomy of the cerebral manifold and CT measures have shown to be strong correlates of cognition (Dickerson et al. 2008). CT changes drastically throughout the healthy lifespan (Salat et al. 2004), and adopts specific patterns in neuropsychiatric disorders (Dickerson et al. 2009; Jubault et al. 2011; Cannon et al. 2015; Bedford et al. 2020). Amongst the most important factors that influence cortical thickness across the lifespan is biological sex (Sowell et al. 2007; Ritchie et al. 2018). Thus, there is an urgent need to understand sex-specific factors that influence brain anatomy through middle age and in the later portions of the lifespan.

Not only are there sex-differences in brain anatomy observed in the later part of the lifespan, but behavioural differences are also well documented. For instance, there is an sex-specific decline in episodic memory and visuospatial abilities (Ferreira et al. 2014). Furthemore, common neurodegenerative disorders, such as Alzheimer’s Disease (AD) and Parkinson’s Disease, have significant differences is sex-specific prevelance rates (Gillies et al. 2014; Association 2016). Some of these sex-differences can be atrributed to the overall higher life expectancy in females (Thornton 2019), or sex-specific prevelance in environmental factors, such as higher education levels in elderly males (Schmand et al. 1997; Coffey et al. 1999; Livingston et al. 2017; Subramaniapillai et al. 2021). Other environmental factors impact men and women differently, such as low BMI, which exerts stronger neuroprotective and pro-cognitive effects in females than it does in males (Moser and Pike 2016), despite the males being generally more active (Barha et al. 2019).

Sex hormones also undergo significant variation in mid-to late-life, and have also been related to brain structure and behaviour. For example, 17β-estradiol is involved in multiple aspects of neural function, including signaling and metabolism (Yao et al. 2010; Brinton et al. 2015), and hippocampal function (Gould et al. 1990; Woolley and McEwen 1992). These observations have led to the hypothesis that estradiol exerts a protective effect against neurodegenerative disease (Rahman et al. 2019), raising questions regarding the impact of the large endocrine fluctuations that take place during the female lifetime. In particular, menopause is accompanied by a drastic decrease in the production of this hormone and the postmenopause is therefore marked by estradiol deprivation (Hoyt and Falconi 2015). In contrast, pregnancy is characterized by hormonal fluctuations that include high levels of estradiol (Barth and de Lange 2020)

However, these endocrine events are not acting in isolation. Many of the risk factors for pathological brain ageing are known to interact with pregnancy and menopause. For example, socioeconomic status, education (Wise et al. 2002; Barnes and Yaffe 2011; Schoenaker et al. 2014;Livingston et al. 2017, 2020), obesity (Barnes and Yaffe 2011; Livingston et al. 2017; Zhu, Chung, Pandeya, Dobson, Kuh, et al. 2018), physical activity (Barnes and Yaffe 2011; Livingston et al. 2017;Floud et al. 2020), depression (Barnes and Yaffe 2011; Georgakis, Thomopoulos, et al. 2016; Livingston et al. 2017; Soares 2019), alcohol consumption (Taneri et al. 2016; Livingston et al. 2020) and smoking (Barnes and Yaffe 2011; Livingston et al. 2017; Zhu, Chung, Pandeya, Dobson, Cade, et al. 2018) have been shown to affect the timing of the menopause transition (MT). Factors such as sleep and social contact can correlate with the lifestyle associated with parenthood (Kalmijn 2012; Hagen et al. 2013;Livingston et al. 2017, 2020). Finally, increased alcohol consumption and anxiety are known to worsen the symptoms associated with the MT, particularly hot flashes (Ashford 2004; Freeman et al. 2005;Schilling et al. 2007; Livingston et al. 2020) and sleep disturbances are commonly reported as a symptom in itself (Cray et al. 2012).

Therefore, in this study, we explore the sex differences in the interactions between contributors to brain ageing using multivariate statistics, namely partial least squared (PLS). Specifically, we perform separate multivariate analyses in males and females that assess interactions between lifestyle factors and CT in a cohort of healthy adults, one of which includes female-specific factors related to menopause. To this end, we used PLS correlation to identify patterns of covariance between risk factors and brain anatomy. Our analyses identified different associations of sex-specific lifestyle and reproductive factors and brain regions in males and females.

## Materials and Methods

### Sample

Cross-sectional data was obtained from the Cambridge Centre for Ageing in Neuroscience (Cam-CAN) data repository (Shafto et al. 2014). Cam-CAN contains multi-modal MRI data on approximately 700 healthy adults aged 18 to 87 which is publicly available to help investigators study the neuroanatomical variation involved in healthy ageing (Taylor et al. 2017). We selected the ages of 40 and 70 years for this analysis, as the lower bound of this range is the minimum age for non-premature menopause (Okeke et al. 2013), and the upper bound allows for a similar ratio of premenopausal and postmenopausal women. The lower bound guarantees that none of the participants will undergo premature menopause in the future. 319 participants (158 females) fit within this range. The final sample consisted of 67 females and 57 males who met the standards of the quality control steps outlined below.

### MRI Acquisition

We used T1-weighted (MPRAGE, 1 mm^3^) structural MRI data collected using a 3T Siemens TIM Trio scanner (Shafto et al. 2014) for the analyses described in this manuscript. All scans were manually quality controlled for motion prior to subsequent image processing described below (https://github.com/CoBrALab/documentation/wiki/Motion-Quality-Control-Manual); (see also: (Bedford et al. 2020)). By removing scans that contain motion artifacts, we can avoid biases in morphometric measurements that are increased in specific subsets of the population (Alexander-Bloch et al. 2016;Pardoe et al. 2016).

### Image Processing

The MRI data was preprocessed using the minc-bpipe-library pipeline (https://github.com/CoBrALab/minc-bpipe-library). This pipeline performed N4 correction (Tustison and Gee 2009), linear registration to Montreal Neurological Institute (MNI) space using BestLinReg (Collins et al. 1994; Dadar et al. 2018), standardization of the field of view and orientation of the brain using an affine template space transform of the coordinate system, and extraction of the brain using a Brain Extraction based on nonlocal Segmentation Technique (BEaST) mask (Eskildsen et al. 2012). The outputs from this pipeline were manually quality controlled, in order to assess whether the image was correctly registered into MNI space, inhomogeneities were corrected and the masks were properly defined.

The scans were then input into CIVET 2.1.0, in order to obtain CT measures across ~40,000 vertices on the cortical surface of each hemisphere (http://www.bic.mni.mcgill.ca/ServicesSoftware/CIVET-2-1-0-Table-of-Contents) (Ad-Dab’bagh et al. 2005). These outputs underwent one final step of manual quality control in order to ensure that tissue classification was performed correctly (https://github.com/CoBrALab/documentation/wiki/CIVET-Quality-Control-Guidelines).

### Demographic Data: Common Variables

Demographic and health related data were self-reported by the participants during a structured interview, performed in their home by an interviewer who entered responses on a computerized script (Shafto et al. 2014). These variables include income and age at which full time education was completed, as well as different measures of social contact from which we used the number of people in the household, the frequency of visits by family and friends, and the frequency of contact with relatives. Participants also reported their number of children, the approximate number of cigarettes smoked throughout their lifetime, as well as the frequency of their alcohol consumption.

Physical activity was measured as energy expenditure excluding rest, or the number of calories burned each day, using the European Prospective Investigation into Cancer Study-Norfolk Physical Activity Questionnaire (EPIC-EPAQ2) (Day et al. 1999; Shafto et al. 2014). A portable, battery-operated weighing scale was used to measure participant weight and a portable stadiometer was used to obtain participant height, from which body mass index (BMI) was calculated (Taylor et al. 2017). Sleep quality was assessed with the Pittsburgh Sleep Quality Index (PSQI) and current severity of depressive symptoms was calculated with the Hospital Anxiety and Depression Scale (HADS) (Buysse et al. 1989; Shafto et al. 2014; Stern 2014).

Some variables of interest were excluded from the analyses due to lack of variance in the data (Table 1). These included hypertension and diabetes, which were also self-reported during the interview. Any missing data was replaced with the mean for the appropriate sex.

**Table 1:**
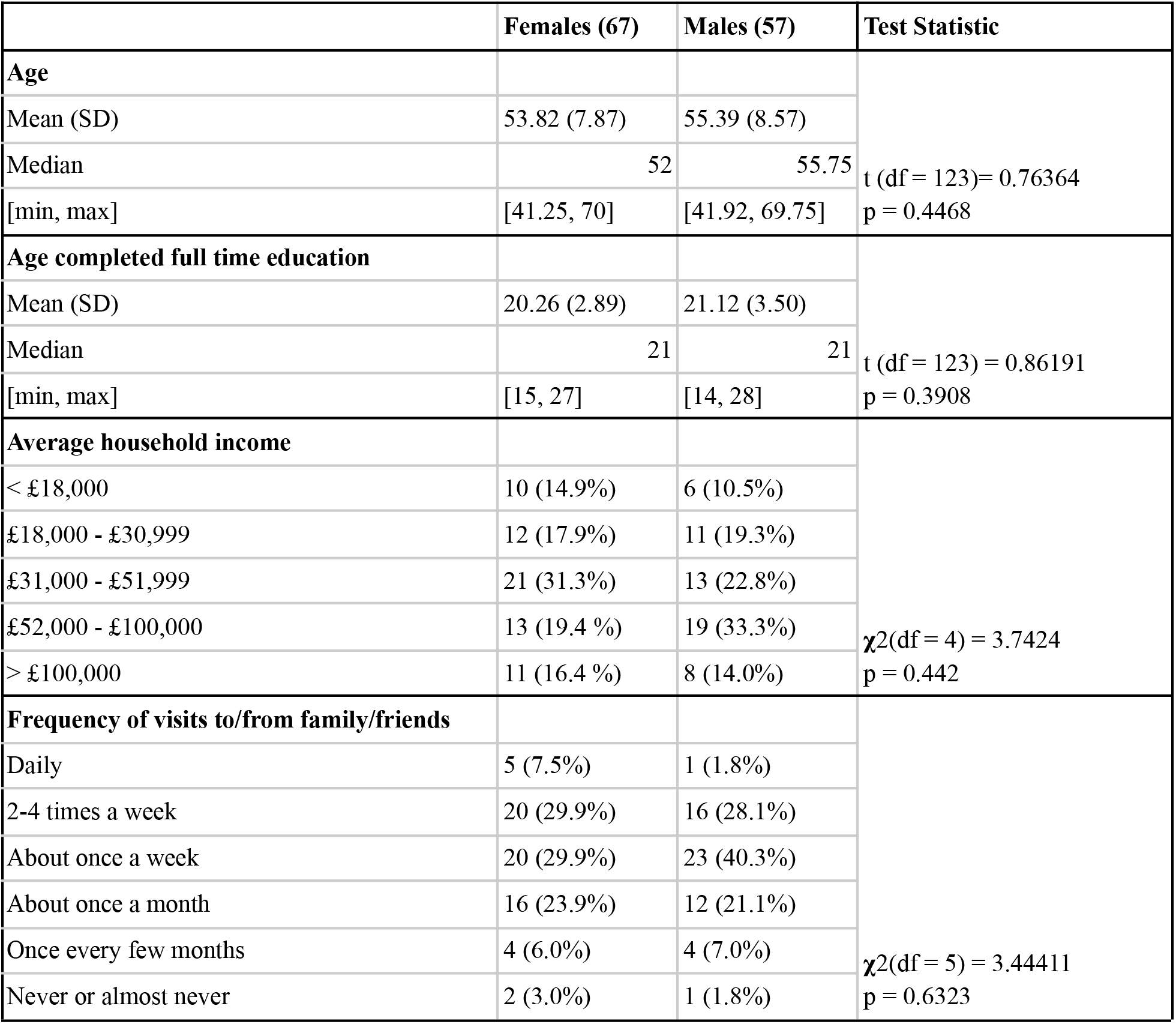
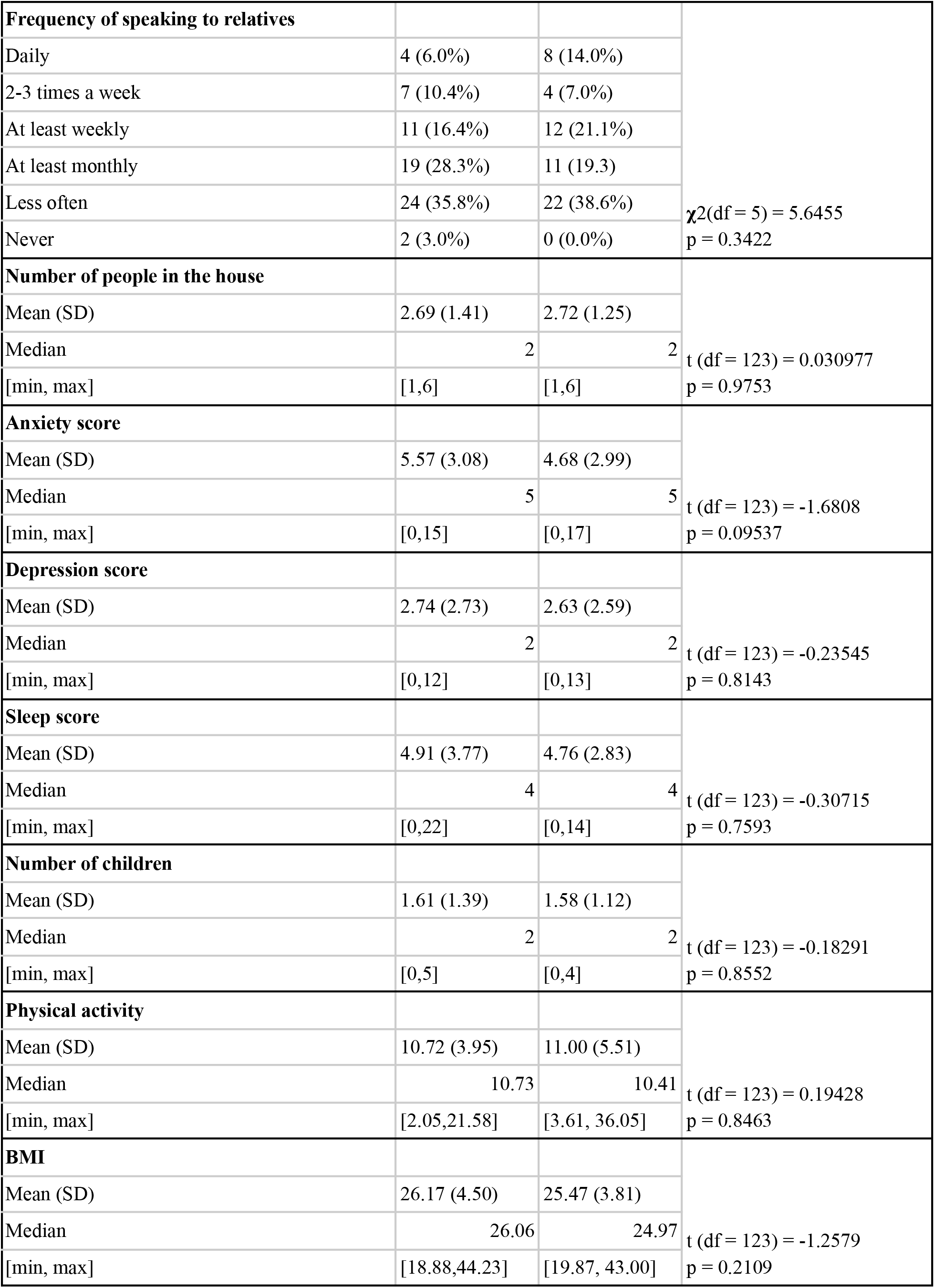
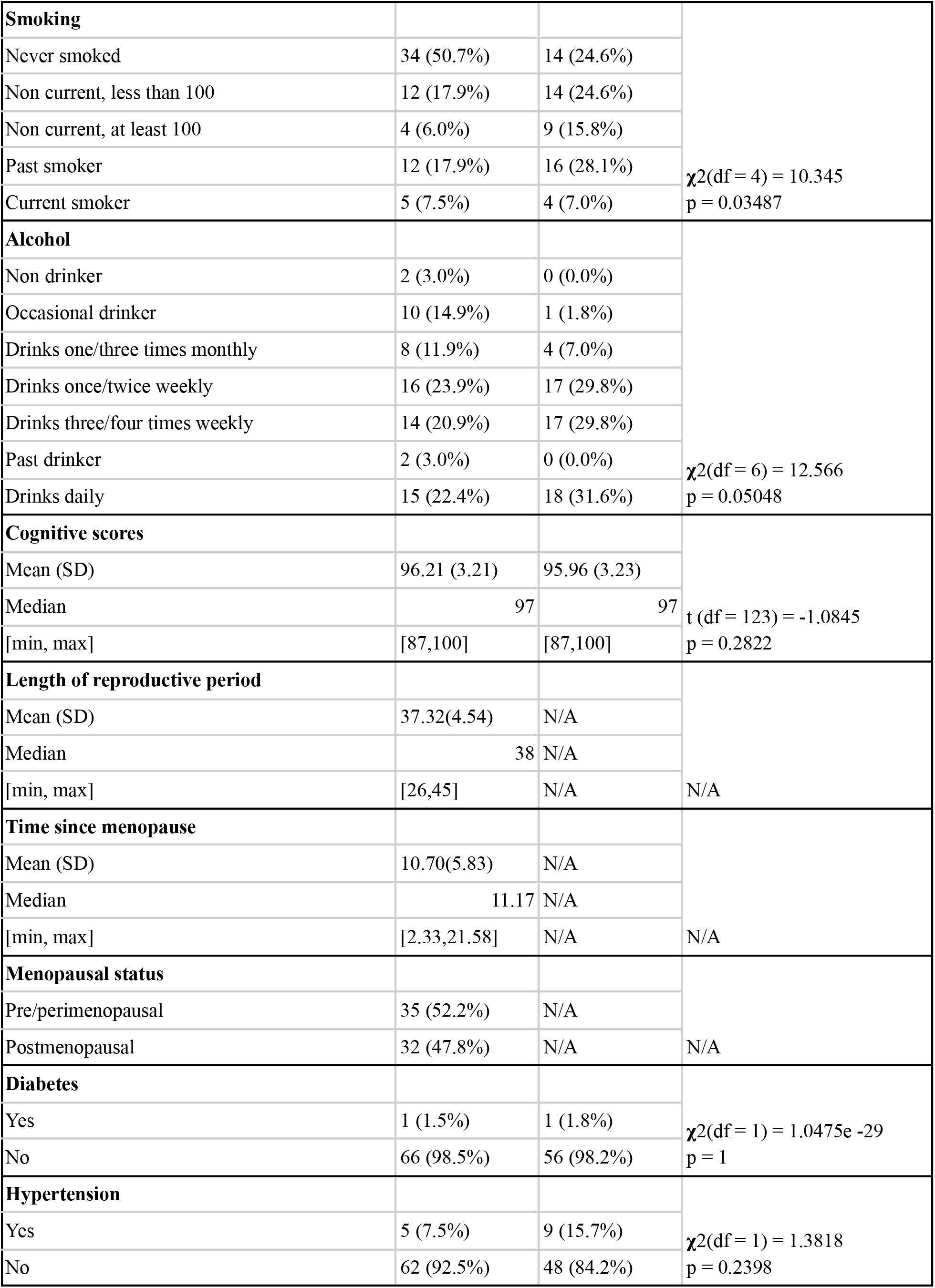

### Demographic Data: Female-Specific Variables

Certain variables that were collected during the same interview were only included in the analyses on females, since there is no equivalent measure in males. These included different measures of reproductive ageing: age at menarche (first period), age at menopause and menopausal status. They were used in order to determine menopausal status, the time elapsed since menopause (zero for premenopausal women) and the duration of the reproductive period, which either corresponds to the number of years between menarche and menopause or between menarche and the time of scan for premenopausal women. Table 1 contains the distribution of menopausal status of all participants as well as of time since menopause and length of the reproductive period of postmenopausal women.

Supplementary table 1 contains the original names and codes of the fields from which the demographic variables were derived.

### Cognitive Data

Cognitive abilities were assessed using the Addenbrooke’s Cognitive Examination Revised (ACE-R) test battery, which evaluates performance across six cognitive domains (orientation, attention, memory, verbal fluency, language and visuospatial ability). In the ACE-R a higher total score signifies better cognitive performance (Mioshi et al. 2006).

### Partial Least Squares

The data was analyzed using partial least squares (PLS) analysis, a multivariate statistical technique that extracts patterns of covariance between two sets of variables (McIntosh and Lobaugh 2004;Krishnan et al. 2011). The analysis was performed using Pyls, an implementation of this technique in Python (v0.0.1; https://github.com/rmarkello/pyls). The inputs to the PLS correlation are two matrices: the brain matrix, which contains cortical thickness values at each of the vertices for all participants, and what is conventionally called the behaviour matrix, which contains the risk factor data for each participant, in the context of our study. The values in both matrices were z-scored along each column prior to performing PLS. The technique consists of a singular value decomposition of the cross-covariance matrix between CT and demographic measures, which, due to z-scoring, this cross-covariance matrix becomes a cross-correlation matrix. The analysis yields a number of latent variables (LV), which are linear combinations of the variables in the two input matrices that maximally covary with each other (Zeighami et al. 2019).

The statistical significance of these LVs is assessed through permutation testing: the rows in the brain matrix are shuffled and the singular value of the latent variable is calculated for each permutation. This creates a null distribution and allows for performance of a null-hypothesis test (Krishnan et al. 2011). Furthermore, the reliability of the contribution of each brain or behaviour variable to a specific LV is calculated using bootstrap resampling. At this step, samples are taken with replacement from the available set of participants and the contribution of each variable to the LV is reassessed. The idea behind this is that if the contribution of a brain or behaviour variable to a LV is highly dependent on which participants are included in the sample, the contribution of the variable is unreliable. This reliability is measured as a bootstrap ratio (BSR), which is calculated by multiplying the weight of the variable in the LV by the singular value of that LV and dividing by the standard error (McIntosh and Lobaugh 2004). A BSR above 3.28 or below −3.28 was considered to be significant and corresponds to a p value of less than 0.001 in a unit normal distribution. Furthemore, the PLS correlation yielded both brain and behaviour scores for each participant, which is a participant-specific measure of the extent to which an individual expresses each of the two patterns that were derived in the analysis (Zeighami et al. 2019).

The PLS analysis were performed separately on males and females, in order to isolate the effect of sex on the interactions between the demographic factors and the brain. Since the focus of this project was the interaction between contributors to brain health during the ageing process and not the process itself, the age variable was regressed out of every demographic and brain variable. This removes the predominant age effects seen in supplementary figures 3 and 4.

### Vertex-wise Linear Models

Vertex-wise general linear models were performed in order to evaluate the individual effect of menopause and the number of children on CT. The models that involved the number of children were performed on males and females separately. Specific information on these models can be found in the supplemental methods.

### Analysis of Cognitive Data

In order to assess whether there was an association between the patterns obtained in the PLS correlation and cognitive performance, general linear models were used to relate brain and behaviour scores with the total score obtained on the ACE-R battery. Since age was residualized from the original data, it was not correlated to the brain or behaviour scores and was therefore not included in the model.

## Results

### Participants

The demographic distribution of the dataset is described in Table 1, which includes the variables that were included in the study, as well as those that were excluded due to lack of variance. Statistically significant differences between males and females were only detected for smoking (p=0.035), where men smoke more than women. The difference between the two groups also tended towards significance for alcohol consumption (p=0.051), where men had higher values.

### PLS in Females

The PLS analysis on the female participants yielded two significant (p < 0.05) LVs. LV1 (Figure 1) explains 47.3% of the covariance (p < 0.001) and demonstrates significant contributions of education, income, length of reproductive period, frequency of contact with relatives and of visits from family to CT in certain regions. These included parts of the left temporal lobe, left prefrontal cortex (PFC) and medial frontal areas as well as some right, frontal regions. LV2 (Figure 2) explained 14.9% of the variance (p = 0.044) and identified increased income, physical activity and alcohol consumption, decreased visits from family and friends and a premenopausal status as significant demographic contributors. These maximally covary with a brain pattern to which CT in the bilateral frontal poles are positively associated and left, medial occipital areas and bilateral somatomotor regions are negatively associated. Thus, both of these LVs show that a combination of variables related to lifestyle and to menopause are associated with CT in frontal regions. An additional PLS was performed in supplementary figure 5, in which the reproductive variables were not included. Supplementary figure 9 contains the same brain maps with a more relaxed threshold of 2.57, or p < 0.01.

**Figure 1:**
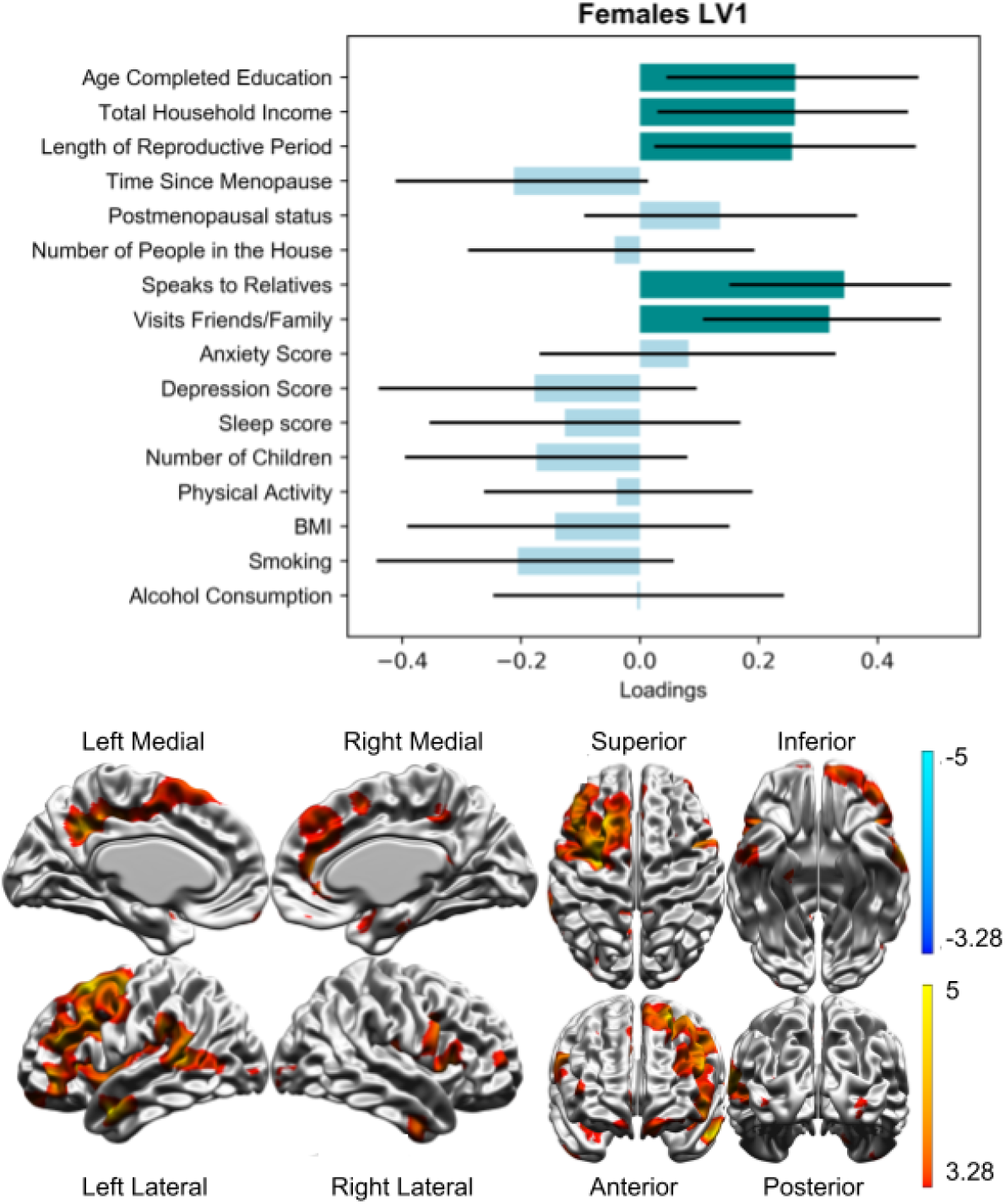
LV1 of the PLS correlation performed on females explained 43.7% of the covariance (p < 0.001). Darker bars on the plot are significant, and maximally covary with the coloured regions on the brain map, where brighter colours indicate increased significance.

**Figure 2:**
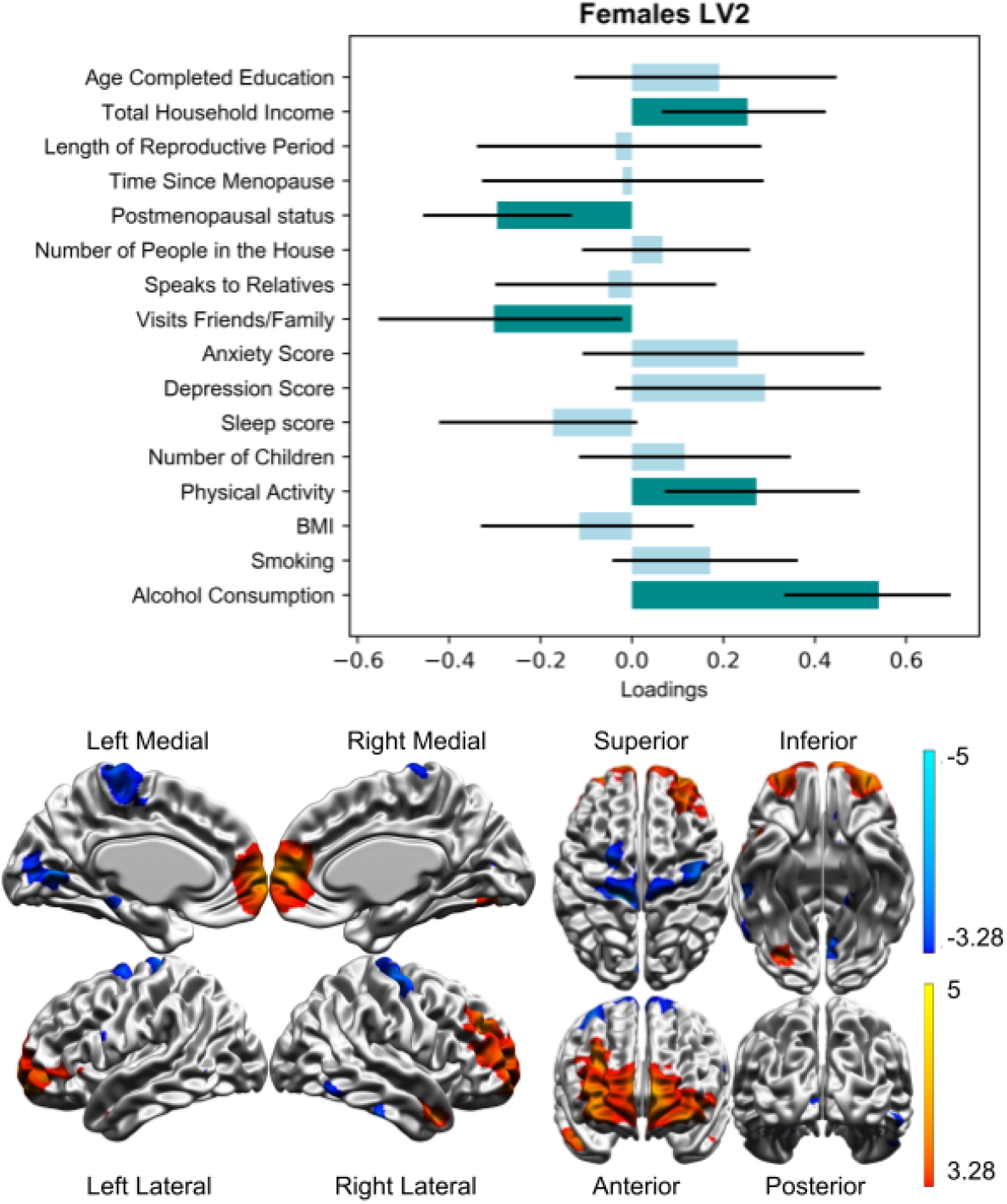
LV2 of the PLS correlation performed on females explained 14.9% of the covariance (p = 0.044). Warm colours on the brain map indicate areas that vary positively with the pattern and cold colours indicate areas that vary negatively with the pattern.

### PLS in Males

In males, the analysis only yielded one significant LV (p = 0.0024) (Figure 3). It explained 48.9% of the variance and associated decreased education, increased anxiety and increased CT in bilateral frontal regions as well as in the medial temporal poles. Some of the small frontal regions identified overlap with those observed in LV1 in females, but do not include the frontal poles, as seen in LV2. The temporal poles, which were found to be significant in males, do not appear in the female LVs.

**Figure 3:**
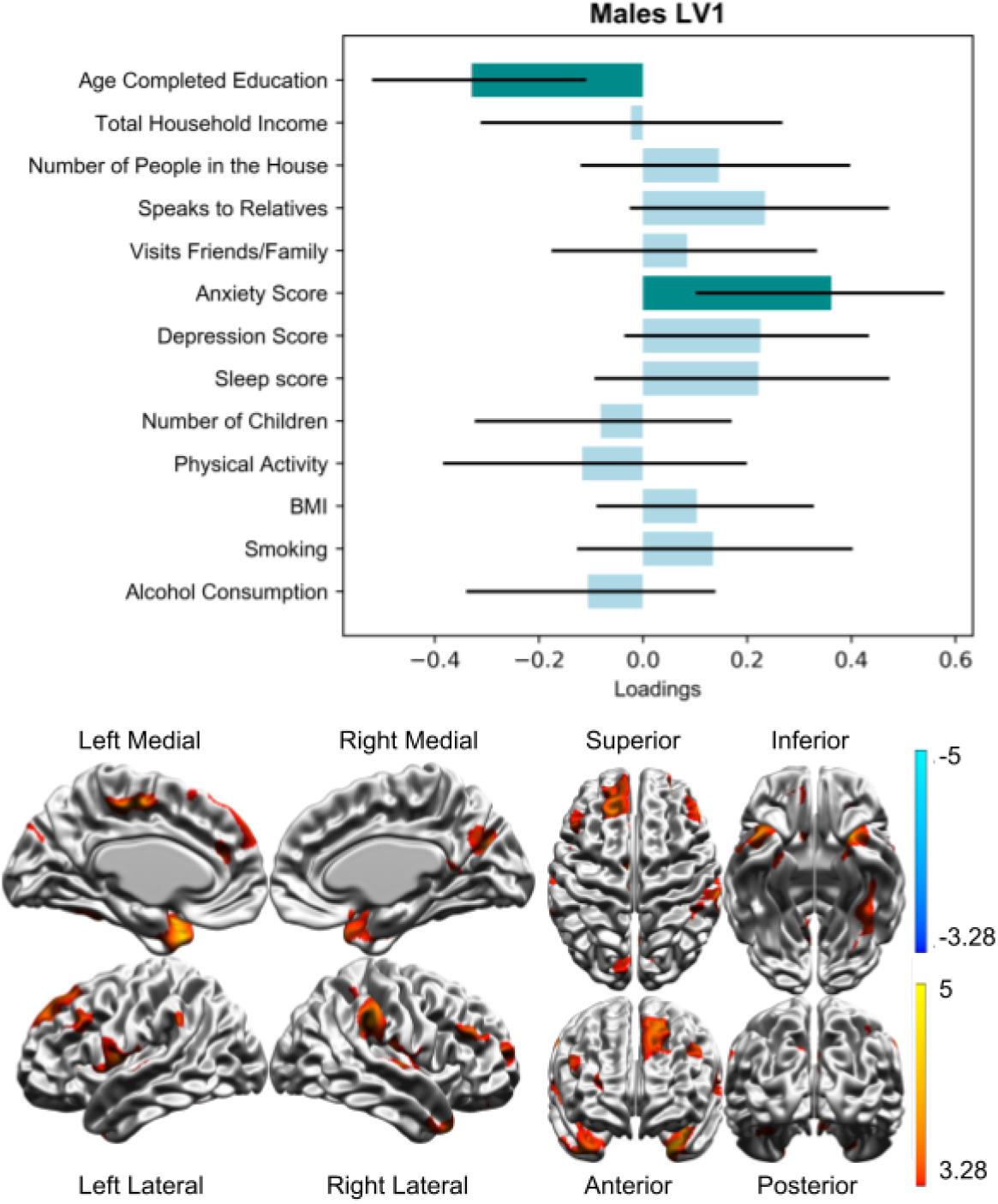
LV1 of the PLS correlation performed on females explained 48.9% of the covariance (p = 0.0024).

### Vertex-wise Linear Models

After correcting for multiple comparisons, no significant associations between menopausal status or number of children and CT were found.

### Cognition

A total of six linear models were used to investigate the relationship between both the brain and behaviour scores corresponding to each of the three LVs described above and the scores obtained on the cognitive tests (Figure 4). These relationships were found to be insignificant both before and after Bonferroni correction.

**Figure 4:**
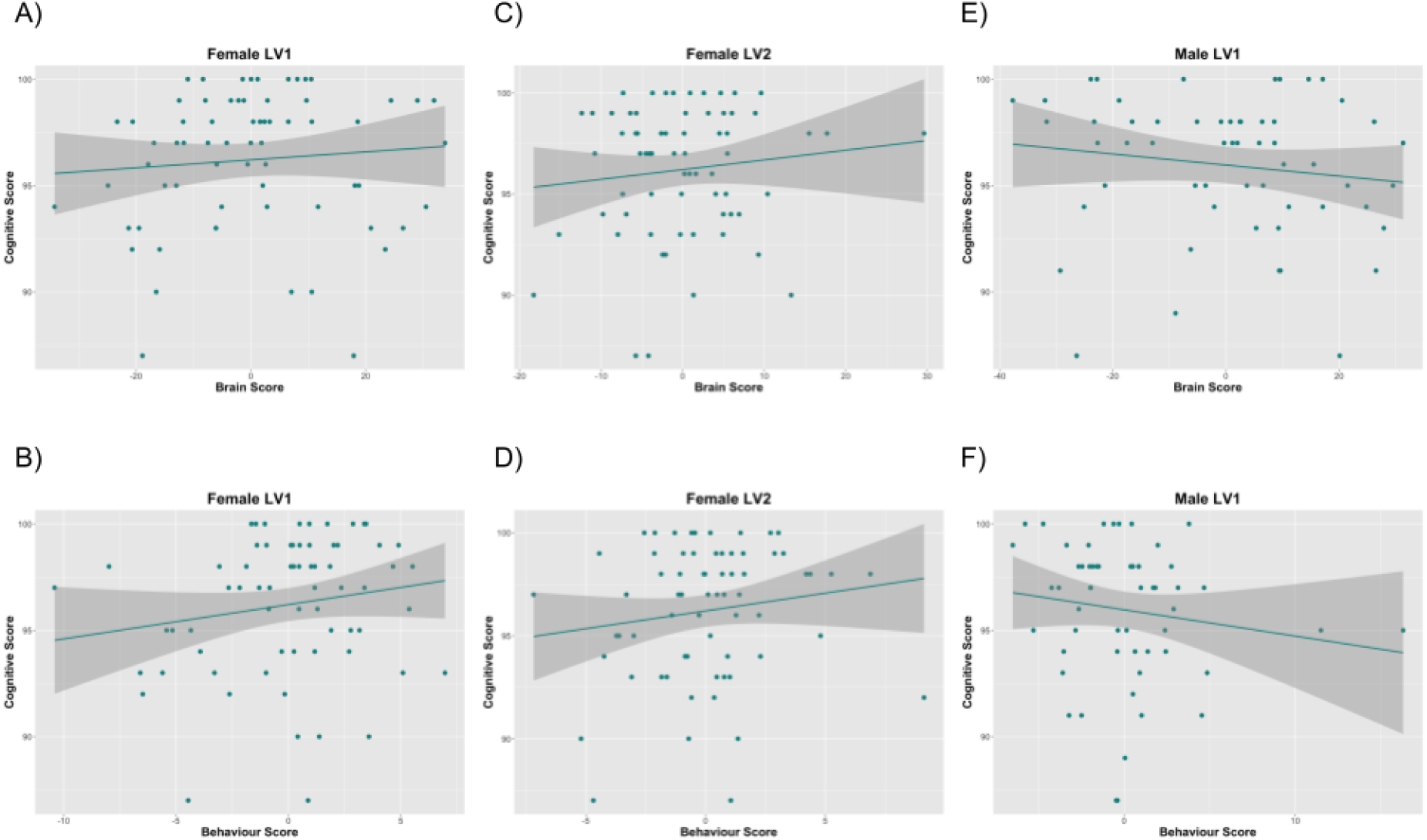
Linear models between cognitive scores and A) brain scores from LV1 in females ( p=0.468, R=0.090) B) behaviour scores from LV1 in females (p=0.162, R=0.173) C) brain scores from LV2 in females (p=0.338, R=0.119) D) behaviour scores from LV2 in females (p=0.214, R=0.154) E) brain scores from LV1 in males (p=0.266, R=-0.150) F) behaviour scores from LV1 in males (p=0.259, R=-0.152)

## Discussion

In this manuscript, we investigate the combined impact of sex, pregnancy, menopause and lifestyle on CT in the context of healthy brain ageing. We did so by performing vertex-wise models, PLS analyses that identified patterns of covariance in the data and linear models that assessed the relationship between these patterns and cognition. While we did not find statistically significant relationships in any of the univariate models, we detected two multivariate association patterns in females and one in males that suggest the lifestyle factors may have a sex-specific impact on brain anatomy. Further, in females there is an indication that this relationship may be driven by interactions with different factors relating to female reproductive health and history.

The PLS analyses performed in each sex yielded very different results. Although both groups had patterns of demographic variables covarying with an increase in CT in the prefrontal cortex and some parietal and temporal regions, the variables driving this increase and the precise cortical regions were entirely different. Additionally, the latent variables that were found to be significant in females included a combination of factors related to both lifestyle and menopause. This indicates that dependencies between the demographic variables driving cortical atrophy in healthy ageing are highly sex-specific and that this effect might be partially driven by interactions between sex hormones and modifyable lifestyle factors.

In all three patterns, we notice that areas that play a role in cognition are positively associated with the LVs. In females, the left ventrolateral prefrontal cortex, left dorsolateral prefrontal cortex and the bilateral frontal poles were identified. These regions play roles in decision making and the cognitive control of memory (Badre and Wagner 2007; Sakagami and Pan 2007), working memory and emotional processing (Levy and Goldman-Rakic 2000; Herrington et al. 2005), and multiple aspects of cognitive control (Mansouri et al. 2015; Bramson et al. 2020), respectively. In males, the temporal poles showed a particularly strong association with the pattern, and they have been associated with semantic knowledge and memory (Levy et al. 2004; Tsapkini et al. 2011). Thus, in males and females, these lifestyle factors are associated with regions that function in distinct cognitive activities. This is in line with previous findings linking the demographic risk factors to cognitive decline and dementia incidence (Barnes and Yaffe 2011; Livingston et al. 2017, 2020), yet our findings did not find an association between these patterns and cognitive performance. This indicates that the neuroanatomical effect of these combined factors can be detected even in the absence of behavioural phenotypes or, potentially, precede the onset of decline. Longitudinal analyses are needed in order to test this hypothesis.

From these results, it can be noted that, although vertex-wise linear models did not identify associations between menopause and CT, variables related to menopause have been found to be statistically significant in both female LVs. This indicates that the MT alone does not cause cortical atrophy, but that it can be potentially associated with patterns of CT decrease in the presence of other risk factors. This confirms our hypothesis that there is a three-way association between this endocrine transition, brain ageing and lifestyle factors. In fact, we observe that variables that correlate with increased estrogen exposure generally combine with neuroprotective variables to be related to an increase in CT, which could be explained through the theory of the healthy cell bias of estrogen action. The theory was originally developed to explain the contrasting results obtained in the study of estrogen interventions to prevent the development of AD when the hormone was administered before versus after menopause. It states that although estrogen exerts a neuroprotective effect on healthy brains, it can further harm those that are already neurologically compromised (Brinton 2008). Our data suggests that this theory could be applied to additional risk factors. The patterns shown in the results indicate that measures of estrogen positively load onto patterns of increased CT and neuroprotective factors, which could be interpreted as estrogen exerting positive effects on healthy cells, not any cells.

More specifically, the risk factors that we found to directly interact with menopause and higher CT in the context of healthy ageing in females are increased education, income, physical activity and social contact. Contrarily, we found that an increased alcohol consumption, in the presence of other factors that included a premenopausal status, covaried with increased CT in certain regions, which contrasts the idea of increased alcohol consumption as a risk factor for dementia (Livingston et al. 2020). Although this could be an artifact of the sample size, it can also be explained by the use of multivariate statistics, which aim at identifying patterns that univariate models cannot capture. In this case, the results indicate a positive impact of alcohol in the presence of other factors, not in the general population. Furthermore, there is evidence from previous studies that subthreshold alcohol consumption, as opposed to alcohol disorders, can increase the age at menopause (Kinney et al. 2006) due to the substance’s pro-estrogenic effects (Gill 2000; Taneri et al. 2016) suggesting a positive impact on endocrine health that might be contributing to this pattern.

Similarly, the results in males show covariance between lower educational attainment and increased CT in the context of high anxiety, which contrasts with the established evidence that presents lower educational attainment as a risk factor for dementia (Livingston et al. 2017). However, it is important to note that lower education is also a risk factor for anxiety disorders (Bjelland et al. 2008) which are known to strong show sex effects (Altemus et al. 2014). Although there is significant heterogeneity in the literature, it has previously been reported that certain anxiety symptoms are associated with increased CT in frontal regions (Ducharme et al. 2014; Szymkowicz et al. 2016). These multiple interactions could once again explain the discrepancy between what univariate models have previously reported and what the present study has found.

On the other hand, many of the lifestyle factors included in the study showed no significant effects in either sex, namely depression, sleep, BMI, smoking and the number of children. The lack of significant results for the latter in either group for both the vertex-wise linear models and the PLS correlations do not indicate a long-term impact of pregnancy nor of parenthood on CT in this cohort.

Finally, we did not find any significant relationships between these demographic or CT patterns and cognitive performance for any of the LVs, which might be influenced by the very high cognitive abilities of the participants and thus the small variance in the data (Table 1). We can therefore state that these patterns of risk factors and cortical atrophy are insufficient to cause a detectable decrease in cognition. Thus, we advocate for further studies to assess the effects of these risk factors on more nuanced definitions of cognition..

To better contextualize our results in females, it is important to understand the previous literature on cognitive decline during the menopause transition. Previous studies have linked it to higher incidence of self-reported memory issues, decline in certain aspects of cognition and increased incidence of AD and dementia (Berent-Spillson et al. 2012; Georgakis, Kalogirou, et al. 2016; Khadilkar and Patil 2019; Rahman et al. 2019). Neuroimaging studies have also correlated postmenopausal status with neurobiological signatures related to AD-risk, including increased amyloid-β deposition and changes to the resting state network (Mosconi et al. 2017; Liu et al. 2021). It is however important to note that evidence for cognitive decline is significantly more robust in women who underwent surgical or premature menopause than in those who underwent spontaneous menopause (Edwards et al. 2019).

Similarly, the large increases in estrogen during pregnancy (Barth and de Lange 2020) have been hypothesized to leave long-lasting effects on the female brain. Studies have found contradictory results relating an increased number of children to both a protective and a detrimental effect on cognition and incidence of dementia (McLay et al. 2003; Deems and Leuner 2020). However, it has recently been hypothesized, based on the observed effects of the number of children on males and the lack of long term effects of incomplete pregnancies on the brain, that it could be the lifestyle related to motherhood and not pregnancy itself that causes neurological changes (Jang et al. 2018; Ning et al. 2020).

In addition to sample size and the potential inaccuracies related to self-report data, there are a few important limitations to this study. First, some risk factors for dementia and maladaptive aging had to be excluded from the analyses due to data availability, including diabetes, hypertension and APOE genotype (Livingston et al. 2017). Diabetes is an endocrine disorder that can develop during menopause due to the changes in hormone production (Paschou et al. 2019) and the risk for this disorder is altered by previous history of estrogen exposure (LeBlanc et al. 2017). Similarly, menopause can be accompanied by an increase in systolic blood pressure (Maas et al. 2009) and this change has been found to be a significant predictor of vasomotor symptom severity (Sadeghi et al. 2012). Finally, APOE-ε4 genotype represents a larger risk of developing AD for women than for men (Altmann et al. 2014; Riedel et al. 2016) and it has even been suggested that apolipoprotein E might be part of the pathway through which estrogen exerts its neuroprotective effects (Struble et al. 2008). Furthermore, the lack of information regarding exogenous estrogen consumption, such as through oral contraceptives or hormone replacement therapy, might be a confounding factor in this study. Finally, due to data availability, no clear distinction could be made between premenopausal and perimenopausal women and, due to sample size, no separate analyses could be performed on premenopausal and postmenopausal women.

Ultimately, the present study identified patterns of covariance between sex, lifestyle factors, reproductive factors and CT in healthy adults. These results revealed sex differences in the interaction between lifestyle risk factors and their impact on the brain, which was partially driven by variables related to menopause. Further studies are required in order to integrate a larger number of variables and evaluate their effects on clinical populations.

