## Supplementary Materials for "Sex Differences in Cortical Morphometry during Ageing: Examining the Interplay between Lifestyle and Reproductive Factors"

**Supplementary methods**:

Linear Models: The first model assessed the relationship between menopausal status and cortical thickness, while correcting for age and for age squared. The second two models looked at the relationship between the number of children and CT and were performed separately in males and in females, while correcting for age and age squared in both sexes and for menopausal status in females. A 5% false discovery rate (FDR) was used to correct for multiple comparisons in these models. The analysis was performed using the RMINC package v.1.5.2.2 (https://github.com/Mouse-Imaging-Centre/RMINC).

**Supplementary Figures**

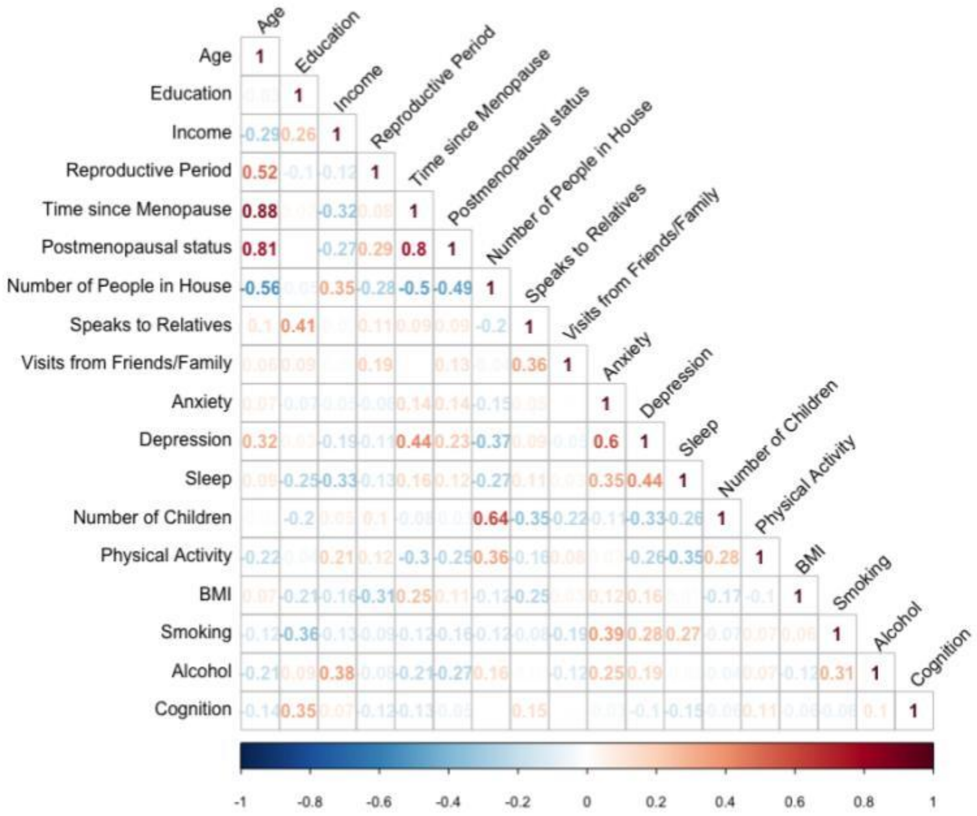


Supplementary Figure 1: Correlation matrix of the demographic variables and the total cognitive score in females. Each number indicates the correlation between two variables, as do the colours, where red represents a positive correlation and blue a negative one. We notice that many of the variables have a very strong relationship with age.

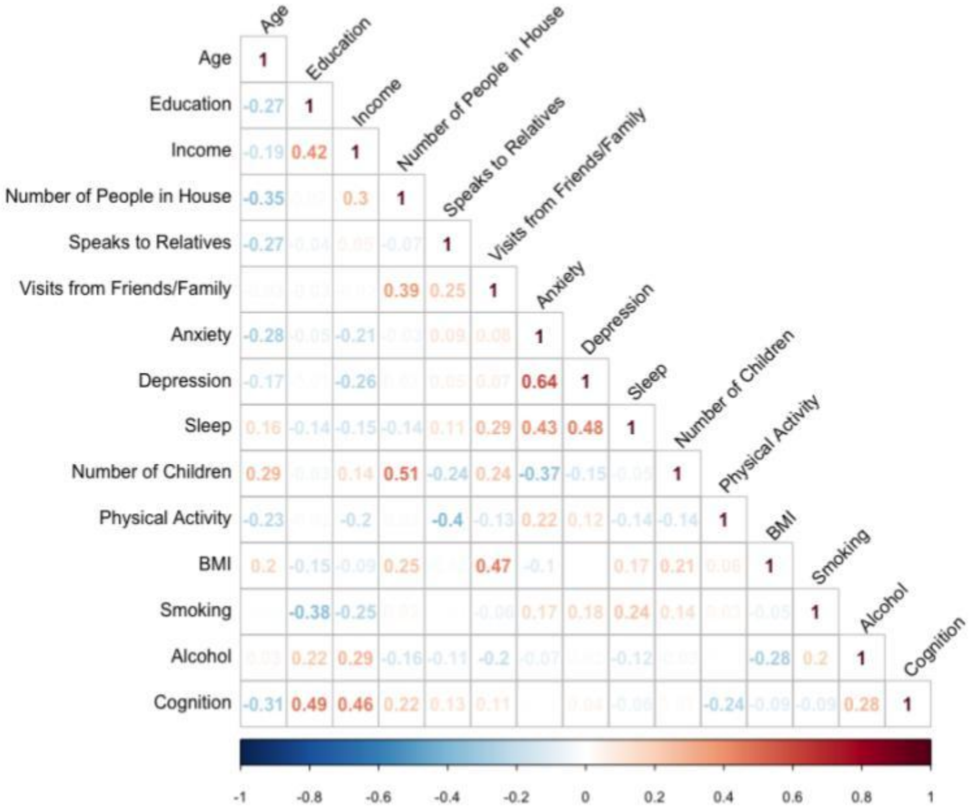


Supplementary Figure 2: Correlation matrix of the demographic variables and the total cognitive score in males.

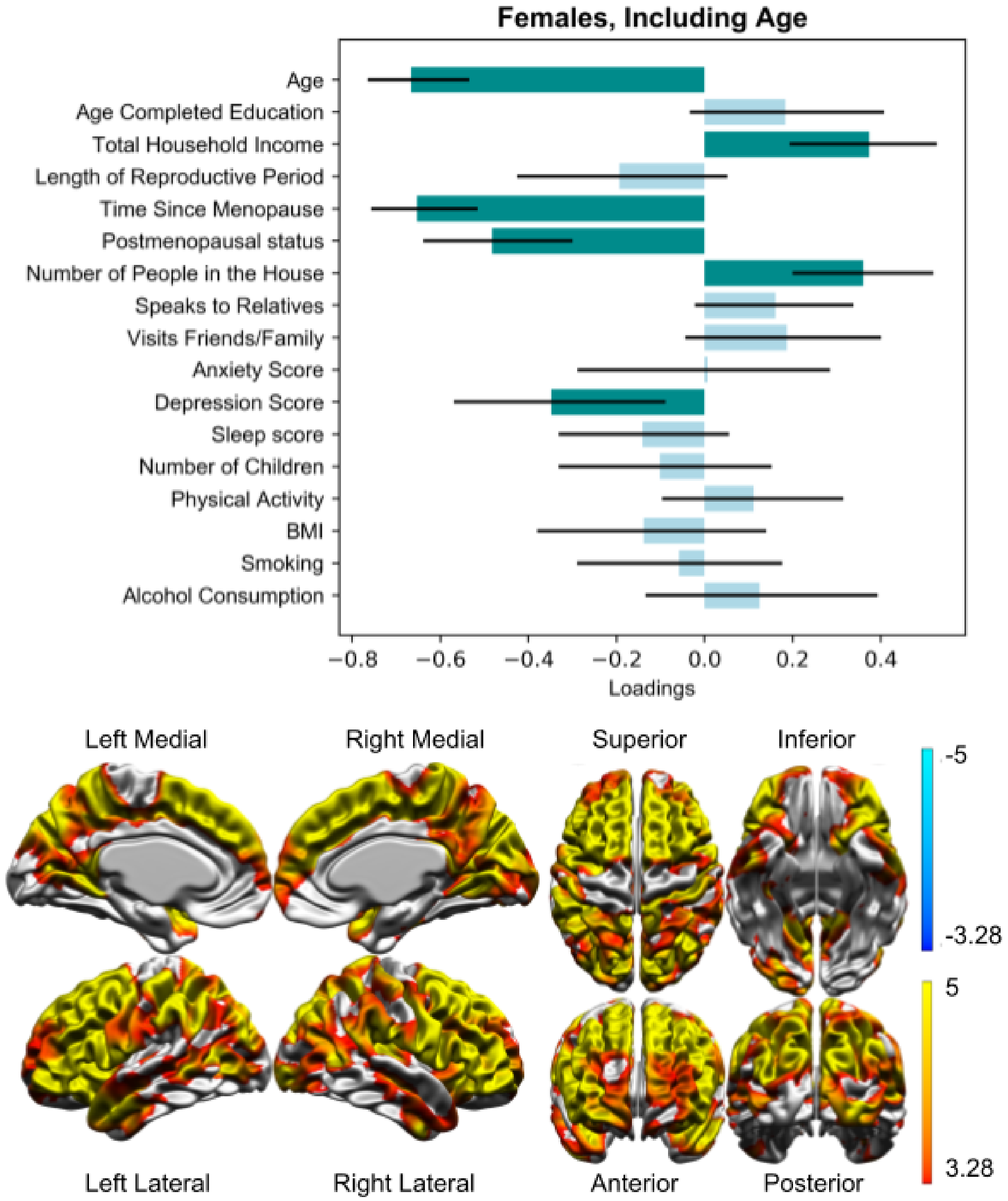


Supplementary Figure 3: PLS results in females where age is included in the demographic variables instead of being regressed out of the data. One LV was found to be significant ( $p<0.001$, 81.8% variance explained) and it shows a very broad increase in CT that covaries with decreased age, time since menopause and depression score, premenopausal status and increased income and number of people in the house. Supplementary figure 1 shows that these five variables are all strong or moderate correlates of age, which implies that this LV mainly shows age effects.

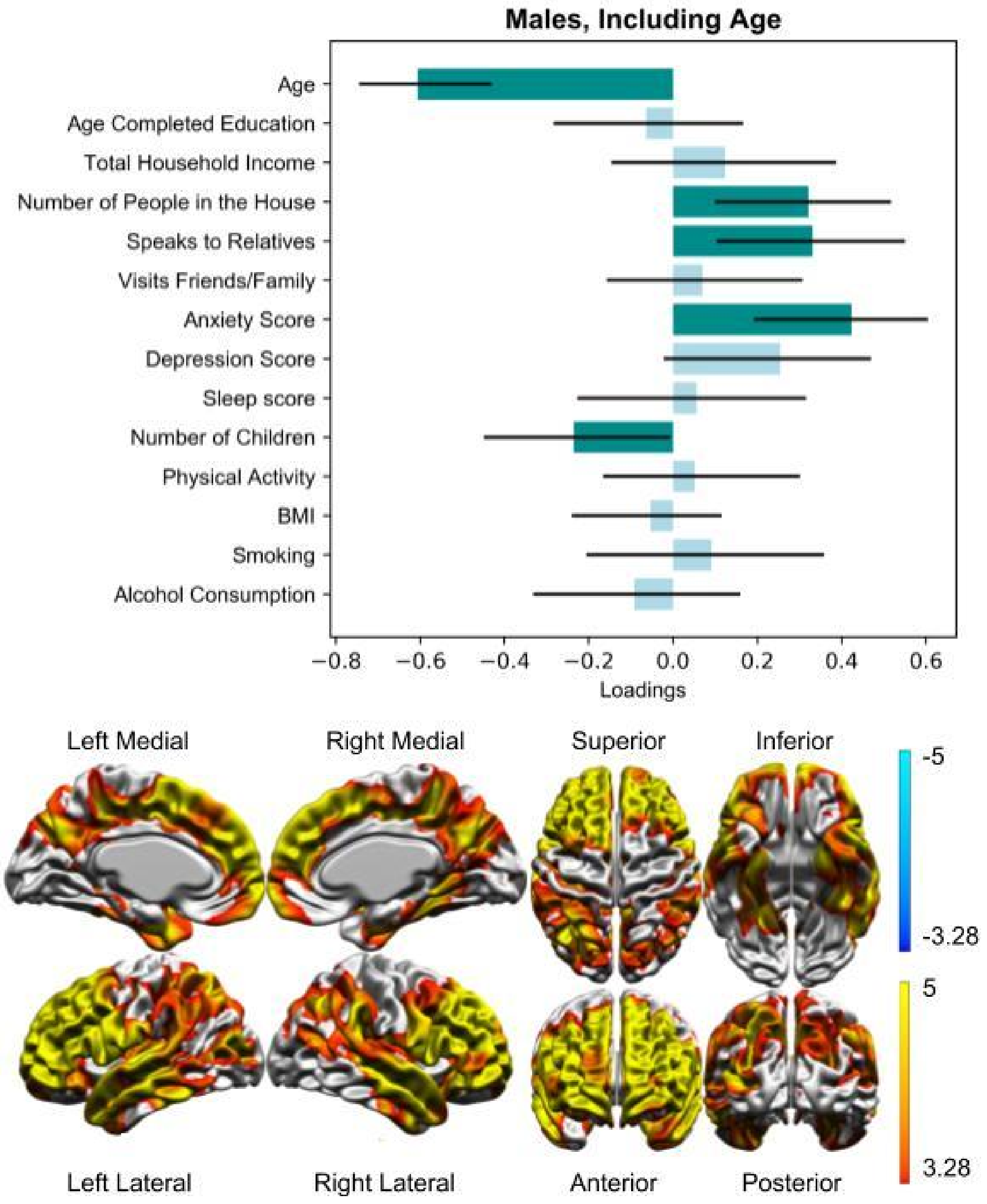


<u>Supplementary Figure 4</u>: PLS results in males when age is included in the demographic variables instead of being regressed out of the data. One LV was found to be significant ( $p < 0.001$, 74.7% variance explained) and, similarly to supplementary figure 3, it shows a very broad increase in CT that covaries with moderate and strong correlates of age, as seen in supplementary figure 2. These demographic variables driving the relationship are the number of people in the house, the frequency of speaking to relatives, anxiety score and the number of children.

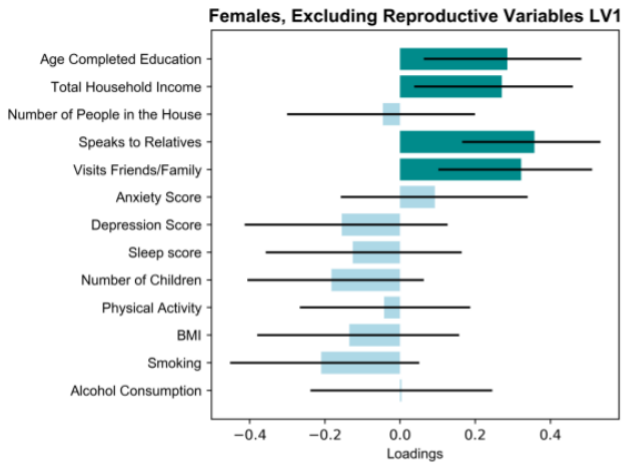
Females, Excluding Reproductive Variables LV1
Age Completed Education
Total Household Income
Number of People in the House
Speaks to Relatives
Visits Friends/Family
Anxiety Score
Depression Score
Sleep score
Number of Children
Physical Activity
BMI
Smoking
Alcohol Consumption
−0.4
−0.2
0.0
0.2
0.4
Loadings

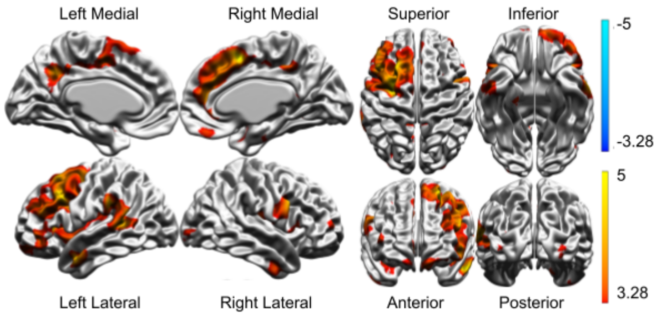
Left Medial
Right Medial
Superior
Inferior
-5
-3.28
5
3.28
Left Lateral
Right Lateral
Anterior
Posterior

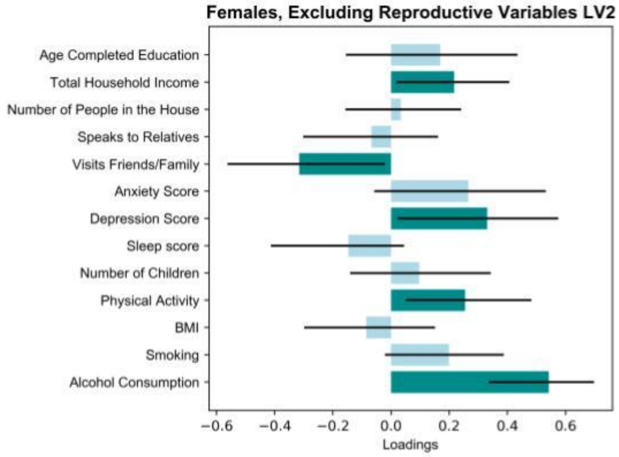
Females, Excluding Reproductive Variables LV2
Age Completed Education
Total Household Income
Number of People in the House
Speaks to Relatives
Visits Friends/Family
Anxiety Score
Depression Score
Sleep score
Number of Children
Physical Activity
BMI
Smoking
Alcohol Consumption
−0.6
−0.4
−0.2
0.0
0.2
0.4
0.6
Loadings

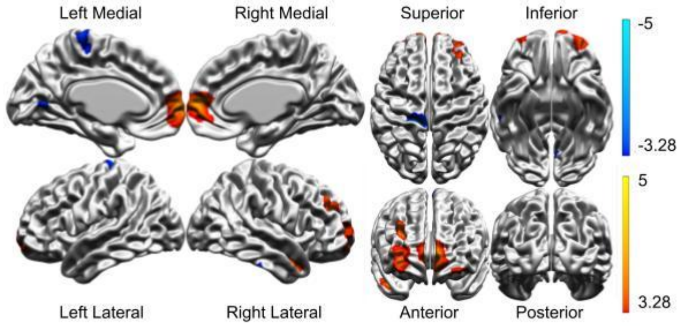
Left Medial
Right Medial
Superior
Inferior
-5
-3.28
5
3.28
Left Lateral
Right Lateral
Anterior
Posterior

Supplementary Figure 5: PLS correlations that excluded the reproductive variables were performed on the female participants in order to allow for a more direct comparison with the results in males. It yielded one significant LV ($p<0.001$, 47.5% variance explained) seen in the supplementary figure 5.A. The brain pattern is very similar to the one found when the reproductive variables were included (Figure 1) and the lifestyle factors that significantly contribute to the LV are the same. LV2 in this PLS decomposition tended towards significance ($p=0.051$, 16.8% variance explained) and shows strong similarities to the LV seen in Figure 2. The same brain regions are identified, although the pattern is slightly broader, and changes in CT in these regions covary with similar sets of lifestyle factors. The primary difference between the two patterns is the contribution of the depression scores, which only becomes significant in the absence of the reproductive variables. Overall, we can conclude that the contrasting results that we observe in males and females are not due to the inclusion of measures related to menopause in one of the analyses, since the patterns are generally preserved once these variables are removed. Furthermore, this indicates that although menopause plays a significant role in the pattern, it is not the primary driver of the LV and the other factors have robust interactions with each other.

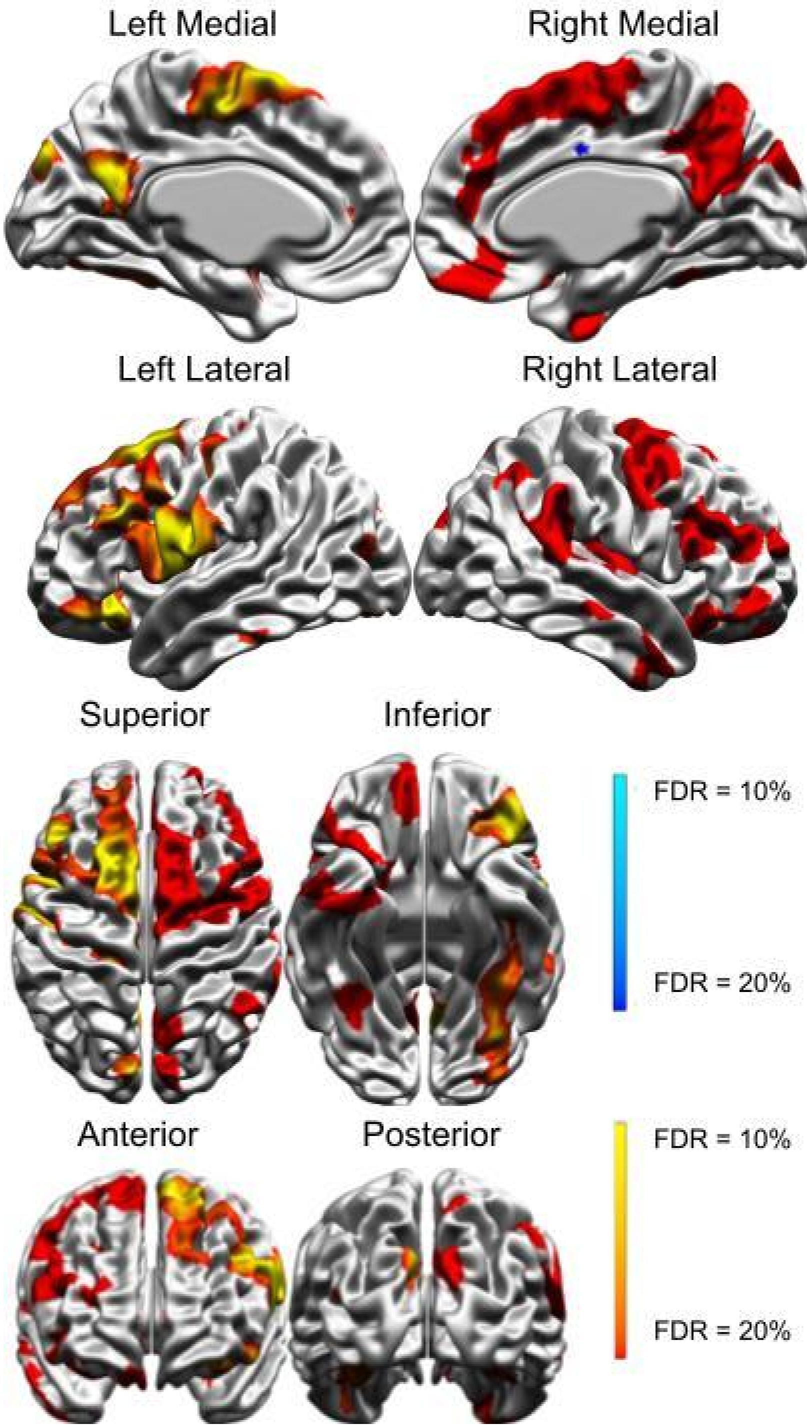


Supplementary Figure 6: Result from a vertex-wise linear model between anxiety and CT in males after correcting for age, age squared, education and income. The results were corrected for multiple comparisons using false discovery rates (FDR) thresholds 20% and 10%. We notice similarities between the regions of the prefrontal cortex identified in this analysis and those in the PLS decomposition on males (Figure 3).

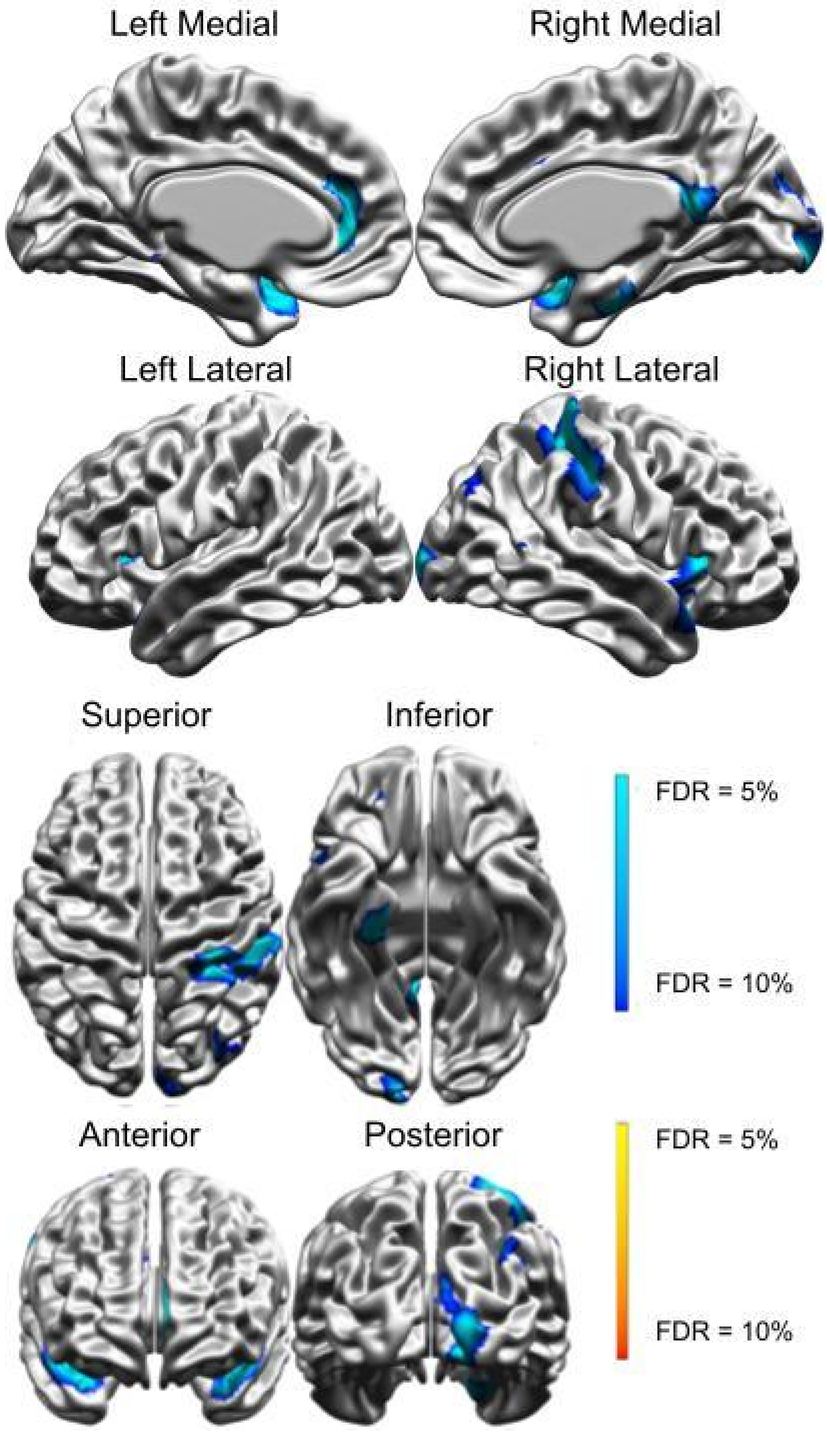


Supplementary Figure 7: Result from a vertex-wise linear model between education and CT in males after correcting for age, age squared and income. The results were corrected for multiple comparisons using FDR thresholds 10% and 5%. The figure shows a decrease in CT in some left, medial frontal and temporal regions and a decrease in CT in right temporal, parietal and occipital areas.

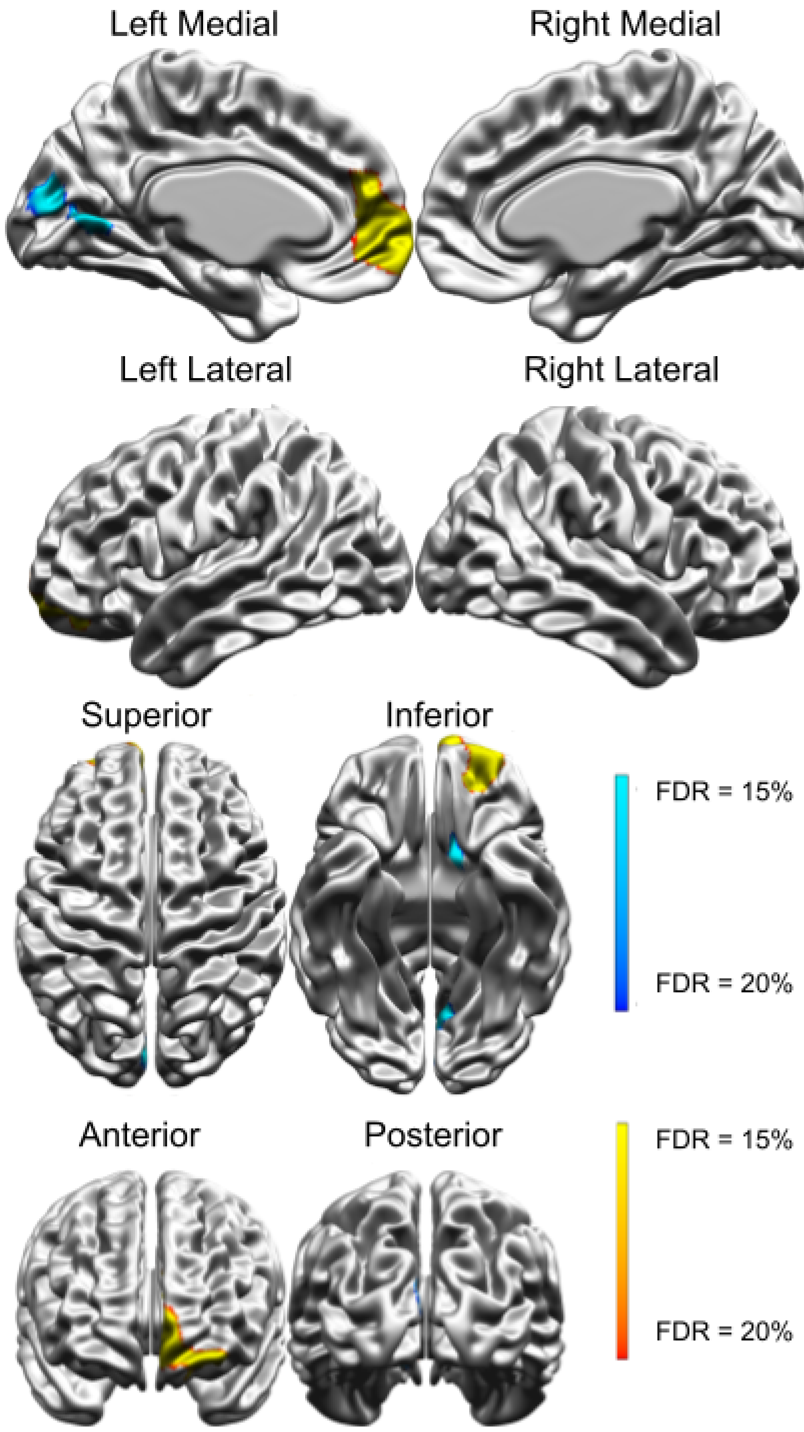


Supplementary Figure 8: Result from a vertex-wise linear model between alcohol and CT in females after correcting for age, age squared, education and income. The results were corrected for multiple comparisons using FDR thresholds 20% and 15%. Similarly to results from LV2 in the PLS analysis on females, we notice a decrease in CT in the medial occipital surface and an increase in CT in the medial frontal pole, yet the pattern is restricted to the left hemisphere.

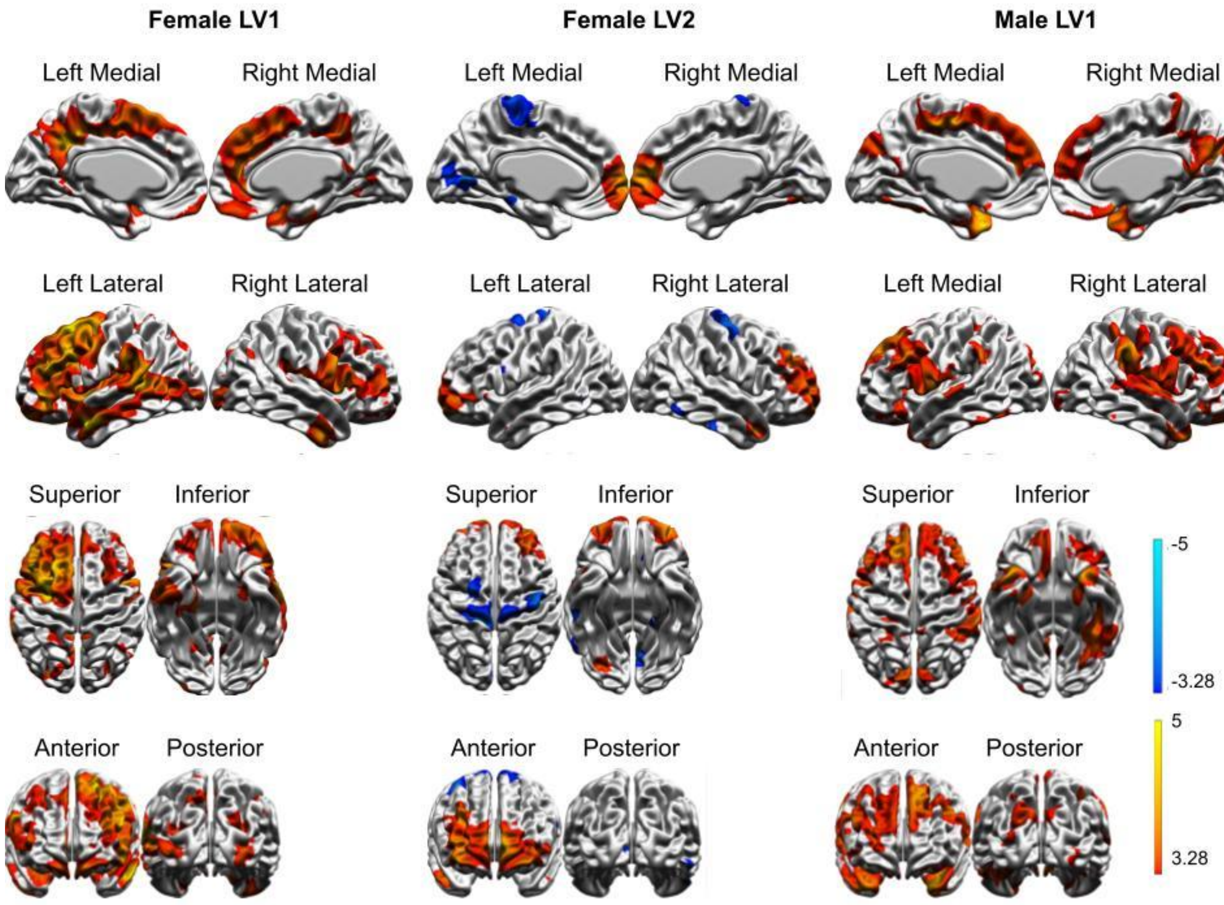


Supplementary Figure 9: Brain maps from Figure 1, Figure 2 and Figure 3 with a threshold of 2.67, which corresponds to $p < 0.01$.

**Supplementary Table:**

| **Variable** | **Original name(s)** | **Code(s)** |
|---|---|---|
| Age | Age | N/A |
| Sex | Sex | N/A |
| Age completed education | Age completed full time education? | homeint_v74 |
| Total household income | Average total household income? | homeint_v15 |
| Length of reproductive period | Age when periods started?, Age when periods stopped? | homeint_v473, homeint_v475 |
| Time since menopause | Age, Age when periods stopped? | homeint_v475 |
| Menopausal status | Had your menopause? | homeint_v474 |
| Number of people in the house | How many people in household? | homeint_v13 |
| Speaks to relatives | How often see relatives to speak to? | homeint_v90 |
| Visits friends/family | How often to (sic) friends family visit you? | homeint_v93 |
| Anxiety score | HADS anxiety score | additional_HADS_anxiety |
| Depression score | HADS depression score | additional_HADS_depression |
| Sleep score | Pittsburg sleep quality index score | additional_psqi |
| Number of children | Number of living children | homeiny_v87 |
| Physical activity | Total energy expenditure discounting rest [net METhrs/d] | epaq_ACTMETS |
| BMI | Weight, Height | physio_cardio |
| Smoking | Smoking summary | additional_smoking |
| Alcohol consumption | Alcohol summary | additional_alcohol |
